# Next generation sequencing analysis of gastric cancer identifies the leukemia inhibitory factor receptor (LIFR) as a driving factor in gastric cancer progression and as a predictor of poor prognosis

**DOI:** 10.1101/2022.05.05.490785

**Authors:** Cristina Di Giorgio, Silvia Marchianò, Elisabetta Marino, Michele Biagioli, Rosalinda Roselli, Martina Bordoni, Rachele Bellini, Ginevra Urbani, Angela Zampella, Eleonora Distrutti, Annibale Donini, Luigina Graziosi, Stefano Fiorucci

## Abstract

Gastric cancer (GC) is the third cause of cancer-related-death worldwide. Nevertheless, because GC screening programs are not cost-effective, most patients receive diagnosis in the advanced stages, when surgical options are limited because the presence of diffuse disease. Peritoneal dissemination occurs in approximately one third of patients with GC and is a strong predictor of poor outcome. Despite the clinical relevance, biological and molecular mechanisms underlying the formation of peritoneal metastasis in GC remain poorly defined. To investigate this point, we conducted a high-throughput sequencing of transcriptome expression in paired samples of normal and neoplastic gastric mucosa in 31 GC patients with or without peritoneal carcinomatosis. The RNAseq analysis led to the discovery of a group of highly upregulated or downregulated genes that were differentially modulated in patients with peritoneal disease in comparison to GC patients without peritoneal involvement. Among these genes the leukemia inhibitory factor receptor (LIFR) and the one cut domain family member (ONECUT)2 were the only two genes that predicted survival at univariate statistical analysis. Because LIFR was the highest regulated gene we have further assessed whether this receptor plays a mechanistic role in GC dissemination. For this purpose, we have first assessed the expression of LIF, a member of IL-6 cytokine family, and LIFR in GC cell lines. Our results demonstrate that exposure of MKN45 cells to LIF, promoted a concentration-dependent proliferation and epithelial-mesenchymal transition (EMT) as shown by modulation of E-cadherin/vimentin gene expression along with JAK and STAT 3 phosphorylation and acquisition of a migratory phenotype. These features were reversed by in vitro treatment with a LIFR antagonist. Together, these data provide support to the notion that development of LIF/LIFR inhibitors might have a role in the treatment of GC.

## Introduction

Gastric adenocarcinoma (GC) is the fifth most common cancer but the third leading cause of cancer-related death (1–3) worldwide (1, 2), with a 5-year survival rate of ≈ 30% (3). The GC is a phenotypically and genotypically heterogeneous disease driven by multiple causative factors, including environmental factors and diet, Helicobacter (H.) pylori or Epstein Barr Virus (EBV) infection and host genetics (4, 5). According to the classical Lauren’s classification, GC is subdivided into three main histological subtypes: diffuse, intestinal and mixed (6, 7). The diffuse subtype is generally more aggressive and predict treatment resistance and poor prognosis (8). In contrast to the diffuse type, the intestinal GC is frequently associated with intestinal metaplasia and H. pylori infection and its prevalence has faced a constant reduction in the last three decades in line with a progressive decrease of P. pylori infection in Western countries (9). While the Lauren histological classification has been widely used over the past decades, its clinical significance remains limited because it does not reflect the molecular heterogeneity of the disease, that has been progressively elucidated by the diffuse application of next generation sequencing technologies to GC (10–13).

The Cancer Genome Atlas (TCGA) and the Asian Cancer Research Group (ACRG) have identified four distinct subtypes of GAC based on genetic and epigenetic signatures: EBV+, microsatellite instability (MSI), genomically stable (GS) and chromosomal instability (CIN) (12, 14). These molecular patterns have been partially validated for clinical use, but there is still a need to better define negative or positive prognostic factors that will predict treatment efficacy.

Currently, radical chirurgical resection is the only therapeutic strategy that offers an effective cure for GC patients (15). However, very often oncologically curative surgery is prevented as most patients are diagnosed in an advanced stage with extensive lymph nodes involvement and distant metastases with limited survival rates. Thus, while Stage IIIC resected tumors associates with 5-year survival rate of 18%, the survival rates for stage IA and IB tumors treated with surgery are 94% and 88%, respectively.

Metastasis is a multistep process(16, 17). A critical event in the formation of metastases is the epithelial-mesenchymal transition (EMT) process in which polarized epithelial cells undergo a process of de-differentiation, characterized by phenotypic changes that are supported by the profound reshaping of EMT biomarkers, including the down-regulation of E-Cadherin, and the upregulation of N-cadherin or vimentin, along with the acquisition of migratory properties (18, 19) and a mesenchymal phenotype. Peritoneal metastases occurs in approximately 30% of patients with GC at the time of diagnosis (20), and their presence impact dramatically on patients survival (21). Furthermore, the peritoneal cavity is a common site of relapse of GC after treatment (22, 23). The poor response of peritoneal carcinomatosis to existing treatments highlights the need to better understand the promoting mechanisms and to identify molecular biomarkers that will predict development of peritoneal carcinomatosis in GC. Recently, NGS studies have shown that the Leukemia Inhibitory factor (LIF) is one of the highest expressed gene in various solid tumors, of stomach (24) (25), pancreas (26), colon (27), liver (28) and breast (29). Of relevance, LIF/LIFR overexpression in these tumours is often associated with a poor prognosis.

LIF is a multirole cytokine that belongs to IL-6 family that plays an essential role in promoting EMT and is envisioned as a potential therapeutic target in many cancers (30). In target cells LIF signaling is mediated by the formation of a heterodimeric complex assembled by the LIFRβ with the glycoprotein (GP) 130 subunit of IL-6 receptor. The GP130 subunit of the receptor is shared with other members of the IL-6 family of cytokines while LIFRβ is shared only with oncostatin M (OSM), cardiotrophin-1 (CT-1), ciliary neurotrophic growth factor (CNTF) and cardiotrophin-like cytokine (CLC). The downstream signaling of the LIF/LIFR pathway involves a JAK-induced STAT3 phosphorylation, AKT and mTor (31).

In addition to its role in directly regulating cancer cells, LIF/LIFR signalling might have a role in promoting the formation of tolerant T cells and macrophages in the tumour microenvironment (32). In this way, LIF has been identified as an important mediator of the immune escape strategies, protecting the tumour from the host’s immune response and promoting growth signals (33). Furthermore, LIF is commonly upregulated in carboplatin and paclitaxel resistant cells, suggesting that it LIF/LIFR overexpression confers chemoresistance to cancer cells (34). Nevertheless, the role of LIF in GC remains unclear and some data suggest that LIF overexpression could be protective (35, 36).

In this paper, we report the transcriptome sequencing (RNA-seq) of paired samples of gastric mucosa and adenocarcinoma samples of GC patients with or without PC and identified LLIFR as one of the highest expressed genes in the GC. LIFR expression is a predictor of PC and poor prognosis. Additionally, by using in vitro cancer cells and pharmacological approaches we demonstrate that inhibition of LIF/LIFR signalling might have utility in the treatment of GC.

## Material and Methods

### Patients and specimens

Gastric carcinoma tissues were obtained from thirty-one patients undergoing surgical resection at the Department of Surgery at the Perugia University Hospital (Italy). None of them received chemotherapy or radiation before surgery. Specimen collection was freshly carried out during surgery by a biologist and paired samples from of normal mucosa sample and neoplastic tissues were collected. Samples were transported to the Gastroenterology laboratory in RNAlater and then snap-frozen at - 80 °C until use. Permission to collect post-surgical samples was granted to Prof. Stefano Fiorucci by the ethical committee of Umbria (CEAS), permit FI00001, n. 2266/2014 granted on February 19, 2014 and by University of Perugia Bioethics Committee, permit FIO0003 n.36348 granted on May 6,2020. An informed written consent was obtained by each patient before surgery.

### AmpliSeq Transcriptome

High-quality RNA was extracted from tumour gastric mucosa and healthy mucosa using the PureLink™ RNA Mini Kit (Thermo Fisher Scientific), according to the manufacturer’s instructions. RNA quality and quantity were assessed with the Qubit® RNA HS Assay Kit and a Qubit 3.0 fluorometer followed by agarose gel electrophoresis. Libraries were generated using the Ion AmpliSeq™ Transcriptome Human Gene Expression Core Panel and Chef-Ready Kit (Thermo Fisher Scientific), according the manufacturer’s instructions. Briefly, 10 ng of RNA was reverse transcribed with SuperScript™ Vilo™ cDNA Synthesis Kit (Thermo Fisher Scientific, Waltham, MA) before library preparation on the Ion Chef™ instrument (Thermo Fisher Scientific, Waltham, MA). The resulting cDNA was amplified to prepare barcoded libraries using the Ion Code™ PCR Plate, and the Ion AmpliSeq™ Transcriptome Mouse Gene Expression Core Panel (Thermo Fisher Scientific, Waltham, MA), Chef-Ready Kit, according to the manufacturer’s instructions. Barcoded libraries were combined to a final concentration of 100 pM, and used to prepare Template-Positive Ion Sphere™ (Thermo Fisher Scientific, Waltham, MA) Particles to load on Ion 540™ Chips, using the Ion 540™ Kit-Chef (Thermo Fisher Scientific, Waltham, MA). Sequencing was performed on an Ion S5™ Sequencer with Torrent Suite™ Software v6 (Thermo Fisher Scientific). The analyses were performed with a range of fold <−2 and >+2 and a p value < 0.05, using Transcriptome Analysis Console Software (version 4.0.2), certified for AmpliSeq analysis (Thermo-Fisher). The transcriptomic data have been deposited as dataset on Mendeley data repository (Mendeley Data, doi: 10.17632/9t86hd78sj.1).

### Statistical analysis

Patients’ descriptive analysis was generated, and their differences were investigated using Student’s t test for quantitative data; normality test accorded to D’Agostino-Pearson was performed, and when not passed, quantitative data were compared using Mann-Withney test. For qualitative data, we used either Fisher’s exact test or Chi-square test. Overall Survival analyses were carried out with Kaplan-Meier’s method, and differences were evaluated using log-rank test. Only variables that achieved statistical significance in the univariate analysis were subsequently evaluated in the multivariate analysis using Cox’s proportional hazard regression model. ROC curves and AUC have also been calculated with the help of statistical software. A p value of less than 0.05 was considered statistically significant. All statistical analyses were performed using the MedCalc Statistical Software version 14.8.1 (MedCalc Soft- ware, Ostend, Belgium), PRISM 7.2 Graph PAD and SPSS, IBM version 23.

### Gastric Cancer cell lines

Human gastric cell lines MKN74, MKN45, KATO III were from the Japanase Collection of Research Bioresources, Human Science Resources Bank (Osaka, Japan). These cells were grown in RPMI 1640 (Sigma-Merk Life Science S.r.l. Milan, Italy) medium supplemented with 10% Fetal Bovine Serum (FBS), 1% L-Glutamine, 1% Penicillin/Streptomycin, in a humidified 5% CO_2_ atmosphere, 37°C. Cells, free from Mycoplasma contamination, confirmed by the use of Mycoplasma PCR Detection (Sigma) were regularly passaged to maintain exponential growth and used from early passages (<10 passages after thawing). To perform all experiments cells were plated, serum starved for 24 h and stimulated for 8- 24-48 h.

### Real-Time PCR

The RNA was extracted from patient biopsies using the Trizol reagent (Invitrogen), and from cell lines using and Direct-zol™ RNA MiniPrep w/ Zymo-Spin™ IIC Columns (Zymo Research, Irvine, CA, USA)., according to the manufacturer’s protocol. After purification from genomic DNA by DNase-I treatment (ThermoFisher Scientific, Waltham, MA USA), 2 µg of RNA from each sample was reverse-transcribed using Kit FastGene Scriptase Basic (Nippon Genetics, Mariaweilerstraße, Düren, Germania) in a 20 μL reaction volume. Finally, 50 ng cDNA were amplified in a 20 μl solution containing 200 nM of each primer and 10 μL of SYBR Select Master Mix (ThermoFisher Scientific). All reactions were performed in triplicate, and the thermal cycling conditions were as follows: 3 min at 95°C, followed by 40 cycles of 95°C for 15 s, 56°C for 20 s and 72°C for 30 s, using a Step One Plus machine (Applied Biosystem). The relative mRNA expression was calculated accordingly to the 2^-ΔCt^ method. Primers used in this study were designed using the PRIMER3 (http://frodo.wi.mit.edu/primer3/) software using the NCBI database. RT-PCR primers used in this study for human sample and human cell lines were as follow (forward (for) and reverse (rev)): CMYC (for TCGGATTCTCTGCTCTCCTC; rev TTTTCCACAGAAACAACATCG), E-CADHERIN (for GAATGACAACAAGCCCGAAT; rev TGAGGATGGTGTAAGCGATG), SNAIL1 (for ACCCACACTGGCGAGAAG; rev TGACATCTGAGTGGGTCTGG), VIMENTIN (for TCAGAGAGAGGAAGCCGAAA; rev ATTCCACTTTGCGTTCAAGG).

### Immunohistochemistry (IHC) and Immunocytochemistry (ICC)

Immunohistochemistry was performed on paraffin embedded human stomach. In brief, Ag retrieval was achieved by incubation of the slides for 90 min in the hot (95 °C) sodium citrate buffer (pH 6.0) and 30 min of cooling at room temperature. Immunostaining technique was carried out using the commercial kit Elabscience®2-step plus Poly-HRP Anti Rabbit/Mouse IgG Detection System (with DAB Solution) (Houston, Texas 77079, USA.) Anti-LIFR Rabbit Polyclonal Antibody (ab235908), Abcam (Cambridge, UK), was incubated overnight at 4°C. Subsequently, sections were incubated with Polyperoxidase-anti-Mouse/Rabbit IgG and then with DAB Working Solution, both supplied by the kit. Slides were counterstained with hematoxylin, dehydrated through ethanol and xylene, and coverslipped using a xylene-based mounting medium.

Slides were observed under microscope and the photos were obtained with the Nikon DS- Ri2 camera, with magnification 20X, 40X, 100X. Immunocytochemistry was performed on MKN45, untreated or treated with LIF 10 ng/mL (14890-H02H, SinoBiological, Düsseldorfer, 65760 Eschborn, Germany). Cells were plate on slides using cytospined. The spots obtained were fixed in 4 % formalin for 20 min and then submitted at the same procedure of immunostaining with the commercial kit Elabscience®2-step plus Poly-HRP Anti Rabbit/Mouse IgG Detection System (with DAB Solution) (Houston, Texas 77079, USA.). After incubation with LIFR primary antibody and secondary antibody supplied by the kit, cells were counterstained with hematoxylin and then observed under microscope with magnification 100x.

### Cell proliferation assay

The cell viability assay was done using the CellTiter 96 Aqueous One Solution Cell Proliferation Assay (Promega, Milano, Italy). It is a colorimetric method for accessing the number of viable cell in proliferation. The MTS assay protocol is based on the reduction of the MTS tetrazolium compound by cells into a coloured formazan product that is soluble in cell culture media. Briefly, on day 0 MKN45 cells were seeded in RPMI 1640 complete medium at 36 *10^3^ cells/100 ul well into 96-well tissue culture plate. On day 1 cells were serum starved for 24 h, and on day 3 cells were primed with LIF (0.5,5,10,50 and 100 ng/ml), or only with vehicle. In another experimental setting, on day 3 cell were pre-treated with EC359 (MedChemExpress, NJ 08852, USA) 25 nM for 1h and finally treated with EC359 () alone or plus LIF, or only with vehicle. After 8-hour incubation period, CellTiter 96 Aqueous One Solution Reagent was added (20 ul/100ul) and incubated until 4 hours at 37°C in a humidified 5% CO_2_ atmosphere. Absorbance was measured using a 96 well reader spectrophotometer (490 nm). Experiments were conducted in tenfold. For analysis the background readings in the wells with medium were subtracted from the sample well read- outs.

### Flow cytometry

The Intracellular flow cytometry staining for Ki-67 was performed using the following Ki-67 Monoclonal Antibody (SolA15), Alexa Fluor™ 488, (eBioscience™, San Diego, California, United States) and DAPI to characterize the cell cycle phases G0-G1, S-G2-M and the Apoptosis rate. Briefly, MK45 cells were seeded in 6-well tissue culture plate (cell density 700 * 103/well) in 100 uL of RPMI 1640 medium supplemented with 10% fetal bovine serum, 1% L-glutamine and 1% penicillin and streptomycin at 37°C and 5% CO2. Cells were serum starved for 24 h and then incubated 48 hours with LIF (10 ng/mL, 50 ng/ml) or only with vehicle. In another experimental setting cells were triggered with LIF 10 ng/ml, EC359 alone or in combination with LIF. Before intracellular IC-FACS staining cells were fixed for 30 minutes in the dark using IC Fixation Buffer (eBioscience™) and then permeabilized using Permeabilization Buffer (10X) (eBioscience™). Flow cytometry analyses were carried out using a 3-laser standard configuration ATTUNE NxT (Life Technologies, Carlsbad, CA). Data was analyzed using FlowJo software (TreeStar) and the gates set using a fluorescence minus-one (FMO) controls strategy. FMO controls are samples that include all conjugated Abs present in the test samples except one. The channel in which the conjugated Ab is missing is the one for which the fluorescence minus one provides a gating control.

### Wound healing assay

MKN45 cells were seeded in RPMI 1640 complete medium at 800*10^3^ cells/well into 24-well plate, so that the day after, they reached above the 70-80% confluence (37). On the Day 1, gently and vertically the cell monolayer was scraped with a new 0.2 ml pipette tip across the centre of the well, during the scratch the medium wasn’t removed to avoid cell death. After scratching, gently the well was washed twice with PBS (Euroclone, Milan, Italy) to remove the detached cells and cell debris, and finally fresh medium contained respectively LIF 10 ng/ml, EC359 100 nM alone or in combination with LIF, was added into each well. Immediately after scratch creation, 24-plate was placed under a phase-contrast microscope and the first image of the scratch was acquired (0h) with OPTIKAM Pro Cool 5 – 4083.CL5 camera. Cells were grown cells for additional 48 h. After 24 h a second image of each scratch was acquired, and after 48 h a third one. The gap distance was being quantitatively evaluated using measuring area. All experiments were performed in triplicate.

### Cell adhesion to peritoneum assay

MKN45 cells were grown in complete RPMI medium; on day 2, cells were starved and left untreated or triggered with LIF (10 ng/ml), EC359 100 nM alone or plus LIF for 48 hours. On day 5, excised parietal peritoneum (∼1.6 cm^2^) was placed in a 24-well culture plate, which had been filled with 1.0 ml of RPMI 1640 medium supplemented with 5% of FBS (38). Gastric cancer cells were detached, fluorescently labelled with BCECF-AM (10 μM) at 37°C for 30 minutes and washed twice with PBS. After trypan blue staining, a suspension of living cells (5 x 10^5^ cells/ml in RPMI 1640) were seeded on the peritoneum in a 24-well plate, and the plate was incubated at 37°C for 60 minutes. After a gentle wash with PBS, the cells adherent to the peritoneum were lysed with 1.0 ml of TRIS 50 mM plus 1% SDS. Fuorescence intensity was measured with a fluorescence spectrophotometer (Ex = 490 nm and Em = 520 nm). Experiments were conducted in quintuplicate.

### Western blot analysis

Total lysates were prepared by homogenization of Raw264.7 in Ripa buffer containing phosphatase and protease inhibitors. Protein extracts were electrophoresed on 12% acrylamide Tris-Glycine gel (Invitrogen), blotted to nitrocellulose membrane, and then incubated overnight with primary Abs against Jak1 (sc-7228 1:500; Santa Cruz Biotechnology), phospho-Jak1 (GTX25493 1:1000; Genetex), STAT3 (sc-8019 1:500; Santa Cruz Biotechnology), phosho-STAT3 (GTX118000 1:1000; Genetex), and GAPDH (bs2188R 1:1000; Bioss antibodies). Primary Abs were detected with the HRP-labeled secondary Abs. Proteins were visualized by Immobilon Western Chemiluminescent Reagent (MilliporeSigma) and iBright Imaging Systems (Invitrogen). Quantitative densitometry analysis was performed using ImageJ software. The degree of JAK1 and STAT3 phosphorylation was calculated as the ratio between the densitometry readings of p-Jak1/ Jak1 and p-STAT3/ STAT3 respectively.

## Results

### Patients

This study includes RNAseq analysis of paired tissue samples from 31 GC patients undergoing surgery in Perugia University Hospital (2013–2019). Peritoneal metastasis dissemination was verified at surgery either macroscopically (P+) or microscopically (Cy+). This led to identification of 19 patients with no peritoneal involvement (P0 and Cy0) and 12 who had peritoneal involvement, (P+ or Cy+) at surgery. Table 1 shows demographic characteristics, primary tumour features and surgical approaches followed in these patients. Patients were then followed up to 5 years after surgery and as shown in Figure 1A median survival time was 41 months and the 5 years overall survival rate was 35.7%. As shown in Figure 1B patients with peritoneal involvement have a significantly worse prognosis, and while patients without peritoneal involvement had a median survival of 53 months (5-year survival rate 49,2%), the median survival time was 14.5 months in patients with peritoneal carcinomatosis (5 years survival rate of 25%).

**Figure 1:**
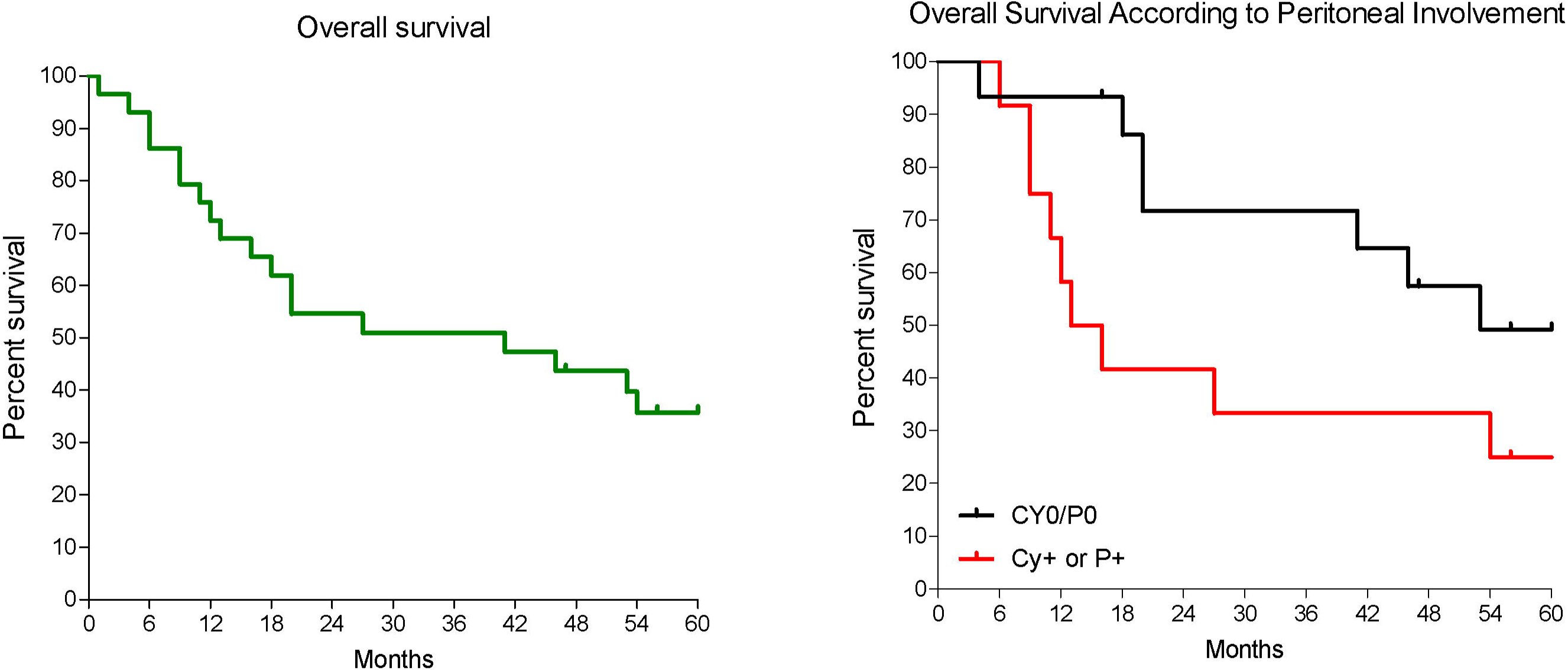
Patients survival. **A)** Overall survival of GC patients and **B)** overall survival GC patients according to the presence of peritoneal disease either macroscopically or microscopically; p < 0.05.

**Table 1.**
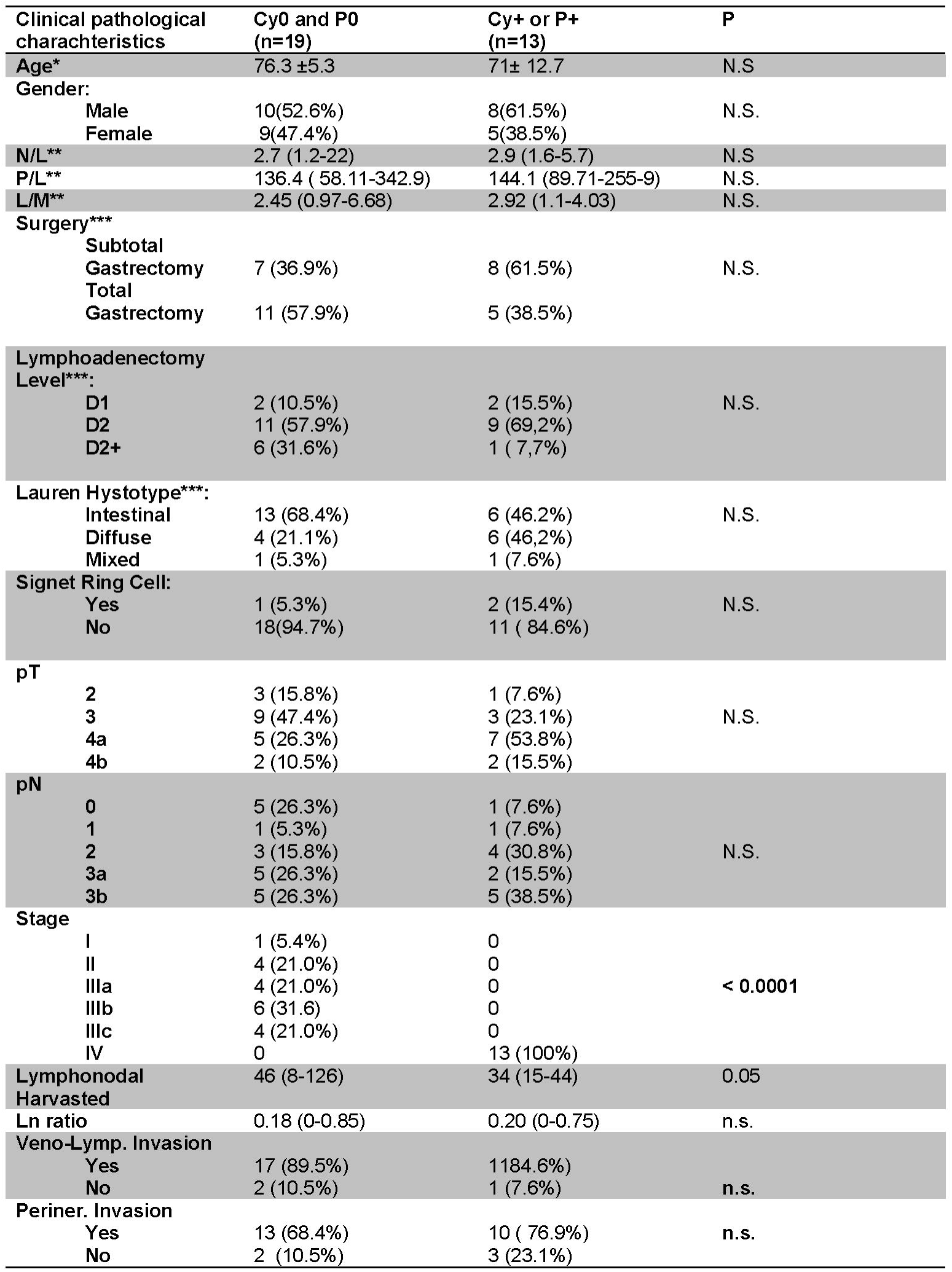
Clinical and pathological characterization of patient population at baseline. Patients were subdivided according to the presence or not of peritoneal disease (Cy0and P0 vs. Cy+ or P+). * Mean and SD; **Median values and range; *** data of Lauren classification were missed in one patient from each group.

### Gene expression profile

The RNA transcription profile by AmpliSeq Transcriptome analysis (RNAseq) of the two patient cohorts, was carried out on paired samples of gastric cancers and their matched normal tissues. The Principal Component Analysis (PCA) of transcriptome shown in Figure2A highlighted the dissimilarities between GC samples obtained from patients with and without peritoneal carcinomatosis, showing only a partial overlap of the two groups. These results were confirmed by Venn Diagram analysis of differentially expressed transcripts. As shown in Figure 2B, this analysis allowed the identification of 341 transcripts belonging to AC+C subsets, that were differentially modulated only in the cancer tissues with patients with peritoneal involvement. Specifically, 79 genes were up-regulated and 262 down-regulated (Figure 2C). The per pathways analysis of these differentially expressed genes using the TAC software (Affymetrix), demonstrated that the most modulated pathway in GC tumoral tissue belong to the EMT pathways, receptors and metabolism, inflammation and signalling clusters (Figure 2D). Analysis of differentially expressed (most upregulated and downregulated genes) in two cohorts of GC patients (with or without peritoneal carcinomatosis) demonstrated that the top three upregulated genes were osteoglycin (ONG), leukemia inhibitory factor receptor (LIFR) and secreted frizzled related protein 2 (SFRP2); while the top three downregulated were, fatty acid binding protein 1 (FABP1), one cut homeobox 2 transcriptional factor (ONECUT2) and Ig superfamily protein glycoprotein A33 (GPA33) genes (Figure 2E). While all six genes showed some degrees of correlation with patient survival (Figure 3) only the relative expression of ONECUT2 and LIFR were statistically correlated with reduced patient survival at univariate analysis (P< 0.05). However, since in comparison to normal mucosa, the expression of ONECUT2 mRNA (39) was upregulated in the bulk tumor of GC patients without peritoneal involvement, but downregulated in those showing peritoneal carcinomatosis, we have focused our attention on LIFR.

**Figure 2:**
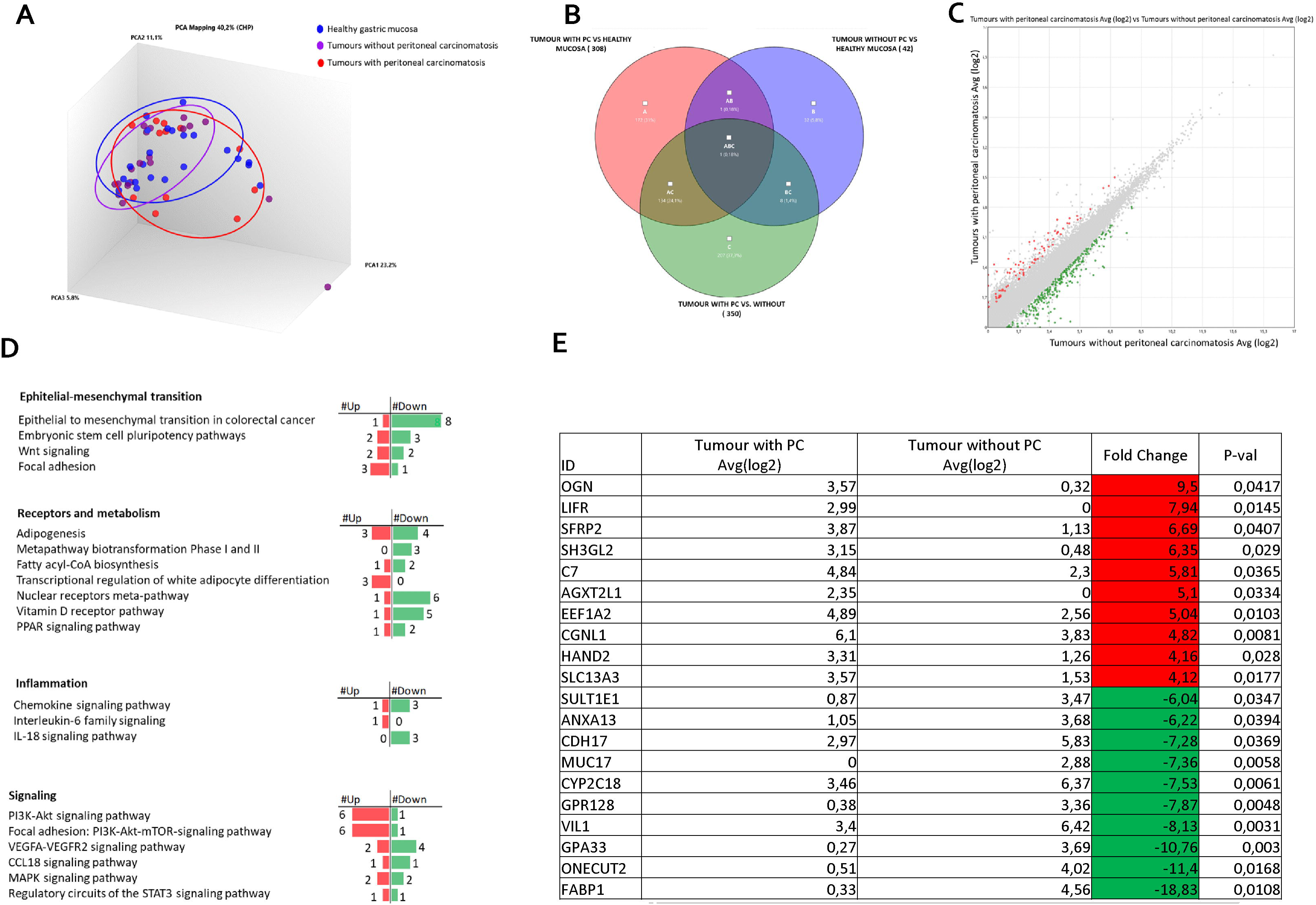
Transcriptome analysis of gastric cancer and paired normal tissues in 31 patients with advanced gastric cancer. **A)** Heterogeneity characterization of gastric samples showed by principal component analysis (PCA) plot. **B)** Venn diagram of differentially expressed genes showing the overlapping regions between the three comparison groups: gastric cancer with peritoneal carcinomatosis vs healthy mucosa (red subset), gastric cancer without peritoneal carcinomatosis vs healthy mucosa (blue subset) and gastric cancer with peritoneal carcinomatosis vs gastric cancer without peritoneal carcinomatosis (green subset). **C)** Scatter plots of transcripts differentially expressed between gastric cancer tissues with peritoneal carcinomatosis and gastric cancer tissues without peritoneal carcinomatosis. **D)** Per pathways analysis of green subset, identification of pathways that can be grouped in four clusters: Epithelial-mesenchymal transition, receptors and metabolism, inflammation and signaling. **E)** Table showing the fold change of expression of the top ten upregulated and downregulated genes included in green subset (Fold Change <−2 or >+2, p value < 0.05).

**Figure 3.**
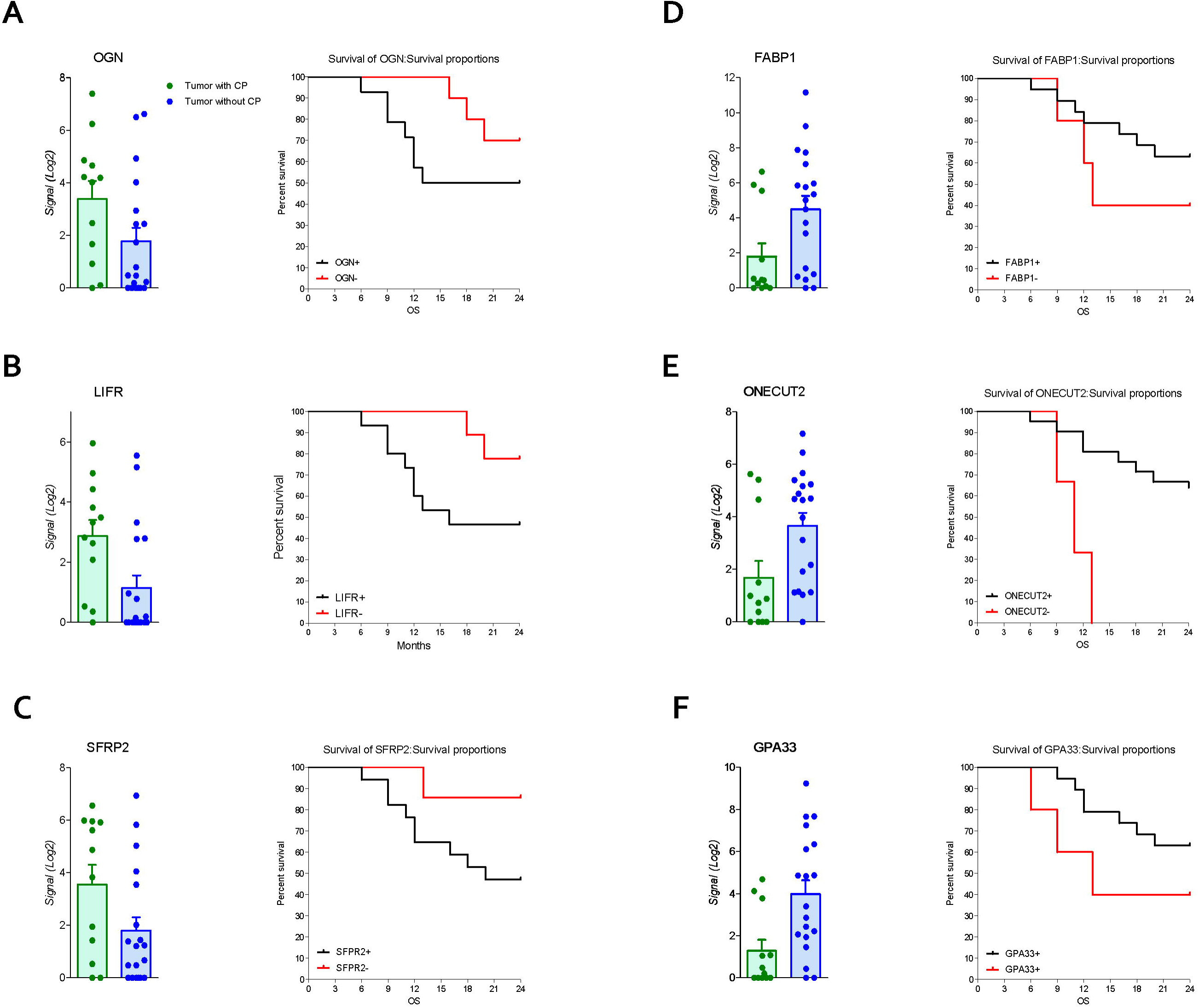
Gene expression and survival curve. Left panel: relative mRNA expression levels extract from RNA-seq analysis and overall survival of patients according to up regulated genes expression of **A)** OGN **B)** LIFR **C)** SFRP2; right panel: relative mRNA expression levels extract from RNA-seq analysis and overall survival of patients according to down regulated genes expression of **D)** FABP1 **E)** ONECUT2 and **F)** GPA33.

### LIFR expression is increased GC patient mucosa with peritoneal carcinomatosis

To explore the role of LIFR and LIF in GC, we have then assessed LIFR expression in 31 tumour samples from GC patients and compared them to the corresponding non-neoplastic mucosa. The results of this experiment demonstrated that LIFR expression in GC tissues was similar to that detected in paired samples of non-neoplastic mucosa (Figure 4A). However, when GC patients with or without peritoneal disease were compared, we found that LIFR expression was significantly increased in patients with peritoneal carcinomatosis (P value <0.05) (Figure 4B). These finding were confirmed by LIFR IHC staining on GC biopsies. As shown in Figure 4, LIFR expression as detected as a faint signal in gastric glands on the normal mucosa, but the signal increased dramtically in the cancer tissues showing a strong localization on the cell membrane of cancer cells (arrow) while some scattered signal was also detected in the tumor matrix (Figure 4F, G). Furthermore, to investigate the role of LIF/LIFR signalling, LIF mRNA expression level was assessed in paired samples of neoplastic and non-neoplastic mucosa of these patients, and as shown in Figure 4C, mRNA LIF expression showed a trend, though not significant, toward reduction in GC samples compared to non-neoplastic mucosa

**Figure 4.**
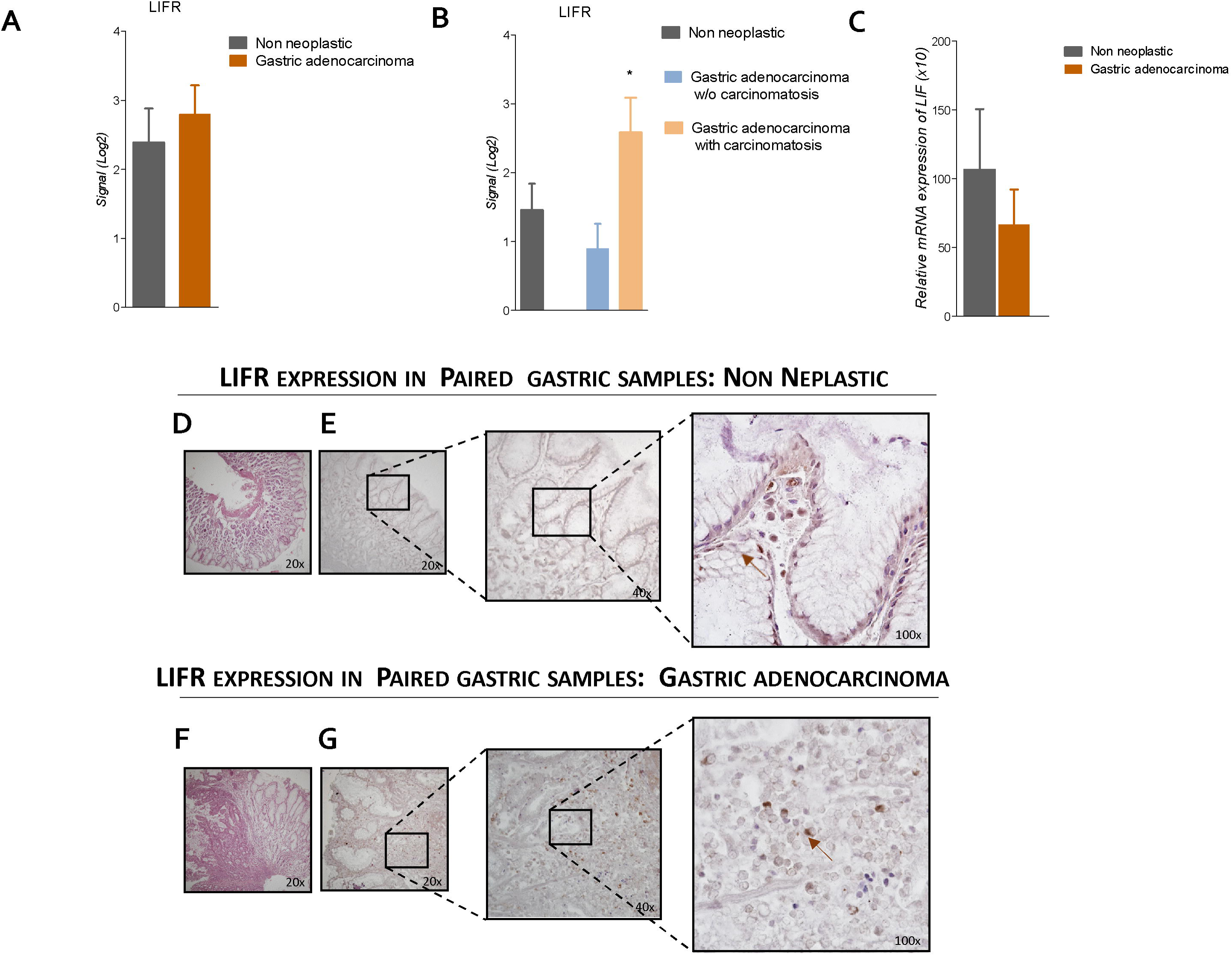
LIFR is a negative prognostic factor for survival of GC patient with carcinomatosis. The expression of LIFR and LIF was examined in surgical samples from non-neoplastic and gastric adenocarcinoma mucosa obtained by GC patients underwent surgery for GC treatment. Data shown are: Gene expression of LIFR (Log2) **A)** in non-neoplastic versus neoplastic mucosa and **B) in** non-neoplastic and gastric adenocarcinoma w/o carcinomatosis versus adenocarcinoma with carcinomatosis. **C)** Relative mRNA expression of LIF. **D)** H&E staining of non-neoplastic mucosa (Magnification 20x). **E)** IHC staining of non-neoplastic mucosa (Magnification 20,40,100x). **F)** H&E staining of gastric adenocarcinoma mucosa (Magnification 20x). **G)** IHC staining of IHC staining of gastric adenocarcinoma mucosa (Magnification 20,40,100x). **H)** Impact of LIFR gene expression on GC patient survival. (*p < 0.05).

### LIF and LIFR expression in GC cell lines

Because the above-mentioned data demonstrated that LIFR expression increases in patients with peritoneal involvement, we have then investigated whether the LIF/LIFR signalling drives the EMT transition using gastric cancer cell lines (Figure 5A, B), and found that the poorly differentiated cell line MKN45 shows the strongest expression of LIFR in comparison to KATO III and the more differentiated cell line MKN74, whose LIFR mRNA expression levels were the lowest. In contrast, LIF mRNA expression displayed an opposite trend, with a lower expression in MKN45 cell and a higher expression by MKN74 (Figure 5A,B), confirming that LIF and LIFR were oppositely regulated as observed in human samples (36) (Figure 4). To further investigate the effect of LIF treatment on GC cells, the MKN45 cell line was selected for further analysis.

**Figure 5.**
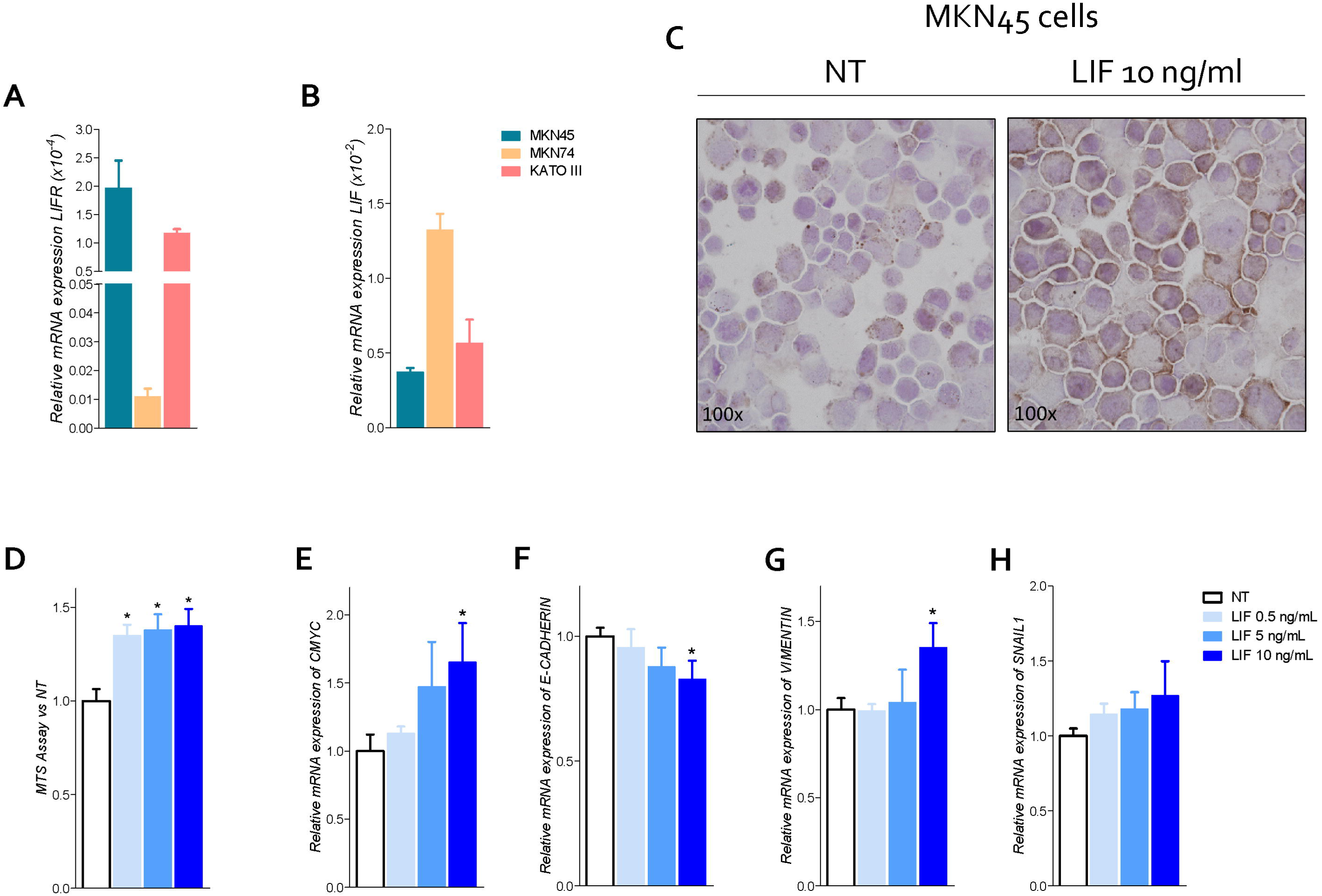
LIFR activation promotes cell proliferation and EMT in MKN45 cells. Relative mRNA expression **A)** LIFR and **B)** LIF in CG cell lines. **C)** IHC staining of LIFR in MNK45 cell lines on left untreated and on right triggered with LIF 10 ng/ml (Magnification 100x). MKN45 cells were serum starved and primed with LIF (0.5,5,10 ng/ml). Data shown are: **D)** Dose-response curve of LIF (0.5,5,10 ng/ml) determined using MTS assay on MKN45 cells. Each value is expressed relative to those of non-treated (NT), which are arbitrarily settled to 1. Results are the mean ± SEM of 10 samples per group. Relative mRNA expression of **E)** the proliferation marker C-MYC; and EMT markers **F)** E-CADHERIN; and **G)** VIMENTIN **H)** SNAL-1. Each value is normalized to GAPDH and is expressed relative to those of positive controls, which are arbitrarily settled to 1. Results are the mean ± SEM of 5 samples per group. (* represents statistical significance versus NT, and # versus LIF, p < 0.05).

To investigate the role of LIF/LIFR in modulating GC cells proliferation and function, MKN45 cells were cultured with increasing concentration of LIF (0.5, 5, 10, 50 and 100 ng/ml). Data shown in Figure 5C demonstrated that LIF induced LIFR expression in LIFR staining cells at ICC (Figure 5C), and also promoted cell proliferation in a concentration-dependent manner as accessed through MTS proliferation assay and relative mRNA expression of CMYC (Figure 5E). Importantly, however challenging MKN45 cells with higher concentrations of LIF, 50 and 100 ng/ml, resulted in a growth-retardation effect, suggesting that at these concentrations LIF might be cytotoxic (S1) (36).

Subsequently, we have investigated the effect of LIF on MKN45 cell cycle and apoptosis (S1). LIF, 10 ng/ml, modulated cell proliferation and cycle, in that the rate of G0-G1 cell was statistically decreased by LIF in comparison to untreated cells, while the rate of cells in S- G2-M phases was increased Figure S1). Again, these effects were biphasic and higher concentrations promoted a cell growth inhibition (Figure S1). Thus, the following experiments were performed using 10 ng/ml LIF as the maximal effective concentration.

Because LIFR promotes EMT in various cell systems, we have investigated the expression of E-cadherin vimentin and SNAIL1, three well recognized biomarkers of EMT in MKN45 cells (40). The results of these experiments demonstrated that exposure of MKN45 to LIF, 0.5, 5, 10, 50 and 100 ng/ml, for 48 h promoted a concentration-dependent reduction in E-cadherin mRNA expression (Figure 5F), which was statistically significative (p< 0.05) at 10 ng / ml, while increased the expression of vimentin and SNAIL1 mRNA in the same range of concentrations (Figure 5 G, H). These effects were lost at higher concentrations, due to increased apoptosis rate and cell growth arrest (Figure S1). Collectively, these data suggest that LIFR agonism promotes cells growth and EMT of GC cells.

To further tight these findings to LIFR activation, we have then investigated whether LIFR inhibition effectively reversed this pattern of regulation. In these studies, we used EC359 a small molecule agent that selectively binds LIFR and down-regulates its pro-oncogenic effects in vitro and in vivo(31). For these purposes, MKN45 were growth in a medium with 10 ng/ml LIF, with or without increasing concentrations, 25, 50,100 and 1000 nM, of EC359 for 48 hours. As shown in Figure 6A, exposure to LIF, again promoted cell proliferation as measured as MST, and this effect was reversed in a concentration-dependent manner by EC359 (Figure 6A), and these effects were statistically significant already at a concentration of 25 nM, while EC359 was cytotoxic at 1000 nM. Similarly, the mRNA expression of CMYC was statistically reduced by 25 nM EC359 (Figure 6 A). Additionally, the LIFR inhibition modulated the cell cycle as shown by Ki-67/DAPI IC-FACS staining (Figure 6D). The cell cycle analysis revealed that EC359 alone did not decreased the rate of proliferative GC cells compared to untreated cells, instead EC359 in combination with LIF effectively reversed the effect of LIF in a statistically significant manner (p<0.05), blocking the shift from resting cell in G0-G1 cell cycle phase, to S-G2-M as also demonstrated by the calculations of ratio between percent of G0-G1 and S-G2-M cells (Figure 6 B). Additionally, EC359 increased the apoptosis cell rates, which was diminished by LIF (Figure 6 C). Consistent with these findings LIFR inhibition by EC359 reversed EMT features in MKN45 cells challenged with LIF. As shown in Figure 6F-H, at the concentration of 25 nM EC359 down-regulated of E-cadherin and reduced the expression of vimentin. Taken together, these data demonstrated that EC359 reversed GC cell proliferation and EMT promoted by LIF/LIFR.

**Figure 6.**
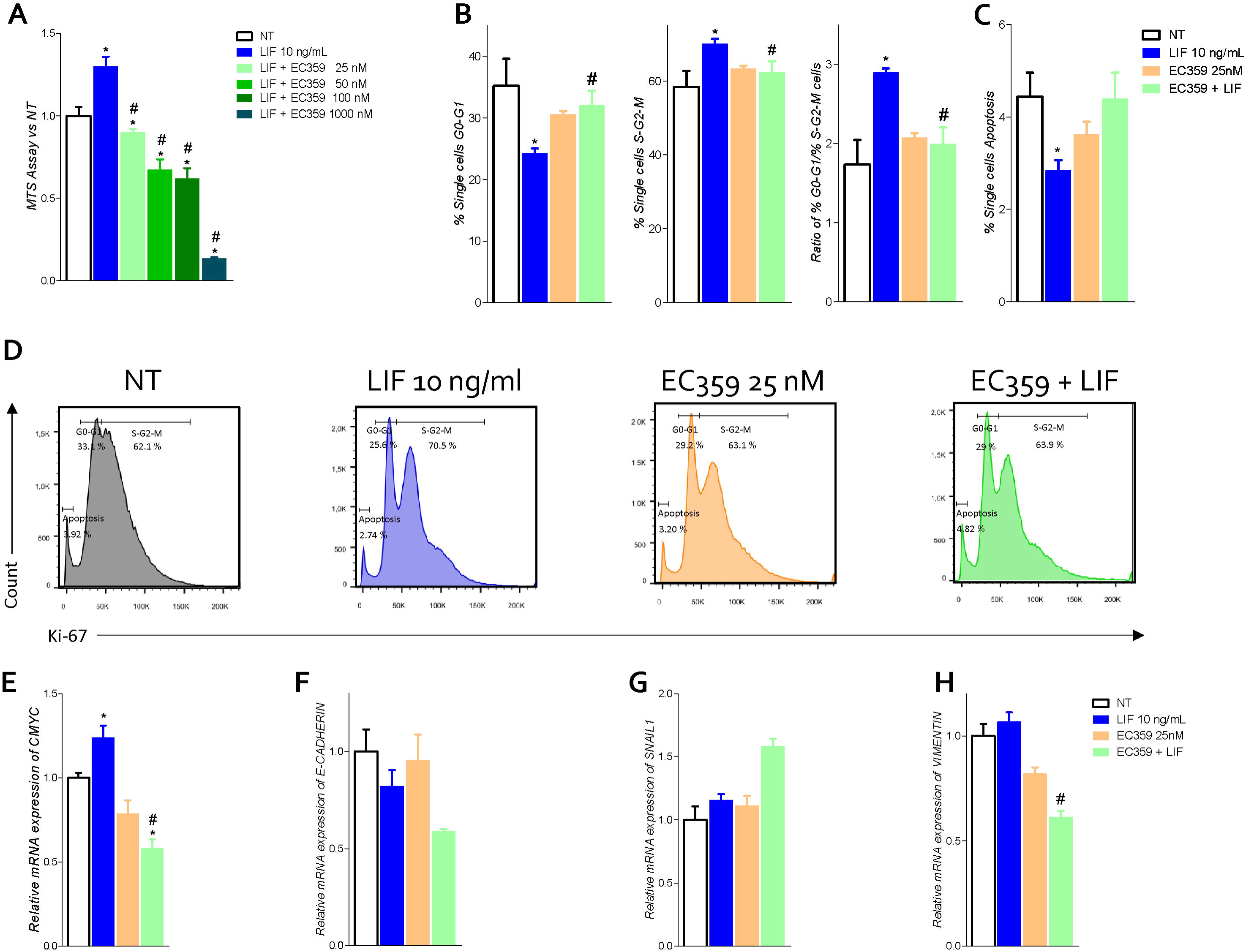
LIFR antagonist EC359 hinder cell cycle progression, increases apoptosis rate in MKN45 cells and inhibits EMT process. **A)** Dose-response curve of EC359 (25, 50,100,1000 nM) determined using MTS assay on MKN45 cells (n=10). MKN45 cells were serum starved and triggered with LIF 10 ng/ml, EC359 100 nM and LIF + EC359 for 48 hours. Cell cycle phases analysis were performed by Ki-67/DAPI staining through IC-FACS. Data shown are: percentage of **B)** from left to right cell in G0-G1 cell cycle phases, S-G2-M cell cycle phases and ratio between % G0-G1 and % S-G2-M; **C)** Percentage of Apoptotic cells. **D)** Representative IC-FACS showed cell cycle fraction and apoptosis rate in NT, LIF 10 ng/ml, EC359 25 nM and LIF plus EC359. Results are the mean ± SEM of 3 samples per group. (* represents statistical significance versus NT, and # versus LIF, p < 0.05). Relative mRNA expression of **E)** the proliferation marker C-MYC; and EMT markers **F)** E- CADHERIN; and **G)** SNAL-1 **H)** VIMENTIN. Each value is normalized to GAPDH and is expressed relative to those of positive controls, which are arbitrarily settled to 1. Results are the mean ± SEM of 5 samples per group. (* represents statistical significance versus NT, and # versus LIF, p < 0.05).

Because LIF/LIFR activation promotes a downstream signalling that involves several kinases, we have then investigated whether challenging MKN45 cells associated with JAK and STAT3 phosphorylation. Western blot analysis reveals that at the concentration of 10 ng/ml LIF increases the expression of LIFR and promotes the phosphorylation of both JAK and STAT3, and that these effects were reversed by LIFR inhibition by 100 nM EC359 suggesting that EC359-mediated inhibitory effect on MKN45 cells involves downregulation of STAT3 signalling (Figure 7A and B).

**Figure 7.**
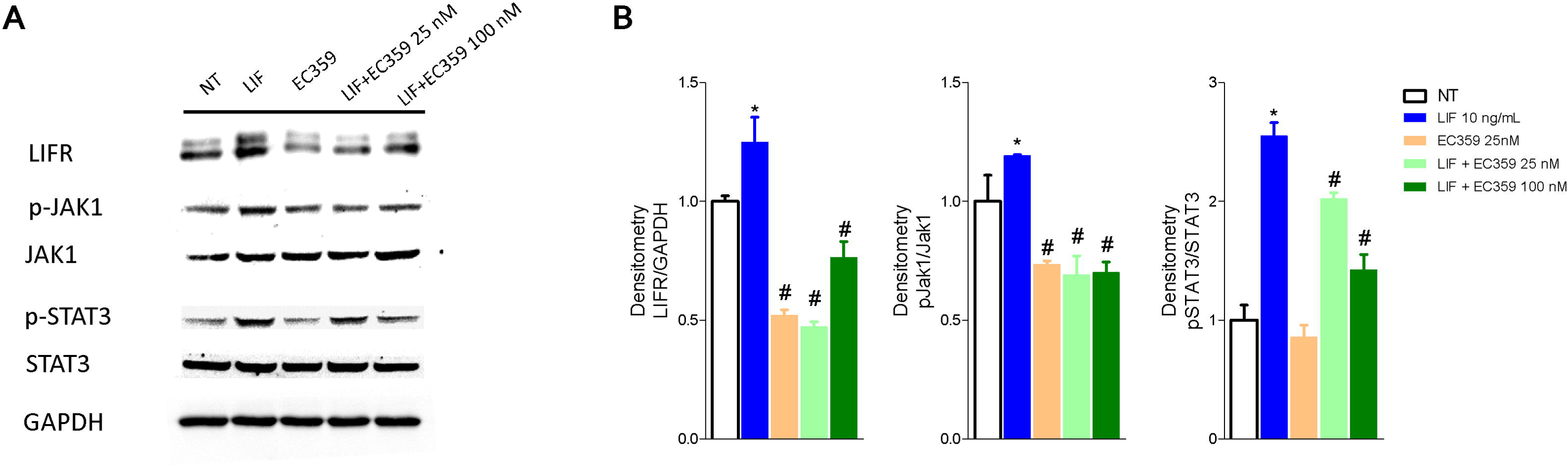
Analysis of JAK-STAT signaling pathway. Representative Western blot analysis of **A)** LIFR, JAK1 and phospho-JAK1, STAT3 and phospho-STAT3, proteins in MKN45 exposed to LIF (10 nM) alone or in combination with EC359 (25 nM and 100 nM) for 20 minutes. GAPDH was used as loading control. **B)** Densitometric analysis demonstrating LIFR expression, phospho-JAK1/JAK1 and phospho-STAT3/STAT3 ratio. The blot shown is representative of another one showing the same pattern.

To evaluated whether modulation of MKN45 by LIF promotes the acquisition of a migratory phenotype, we have performed a scratch wound healing assay, a validated method to functionally assess EMT (Figure 8 A-B). For these purposes MKN45 cells were growth in a complete medium and at the day 0, the scratch was produced as described in Material and Methods and cells migration in response to LIF, 10 ng/ml, EC359, 100 nM, and LIF plus EC359, assessed at different time points, i.e., 0, 24 and 48 h. As illustrated in Figure 8, while exposure to LIF induced a robust wound closure by reduction of the wound area of 45.41 % at 24 hours and 82.23 % at 48 h, this pattern was reversed by exposure to EC359 (p<0.05). Also, EC359 alone reduced the percentage of wound closure compared to untreated cells but these changes were not statistically significant. Similar finding was observed assessing the adhesion of MKN45 to the peritoneum, a model with s translational relevance. In this assay while LIF promoted MKN45 adhesion to the mouse peritoneum the effect was significantly attenuated by co-treating the cells with EC359 by ≈ 30% (Figure 8 C).

**Figure 8.**
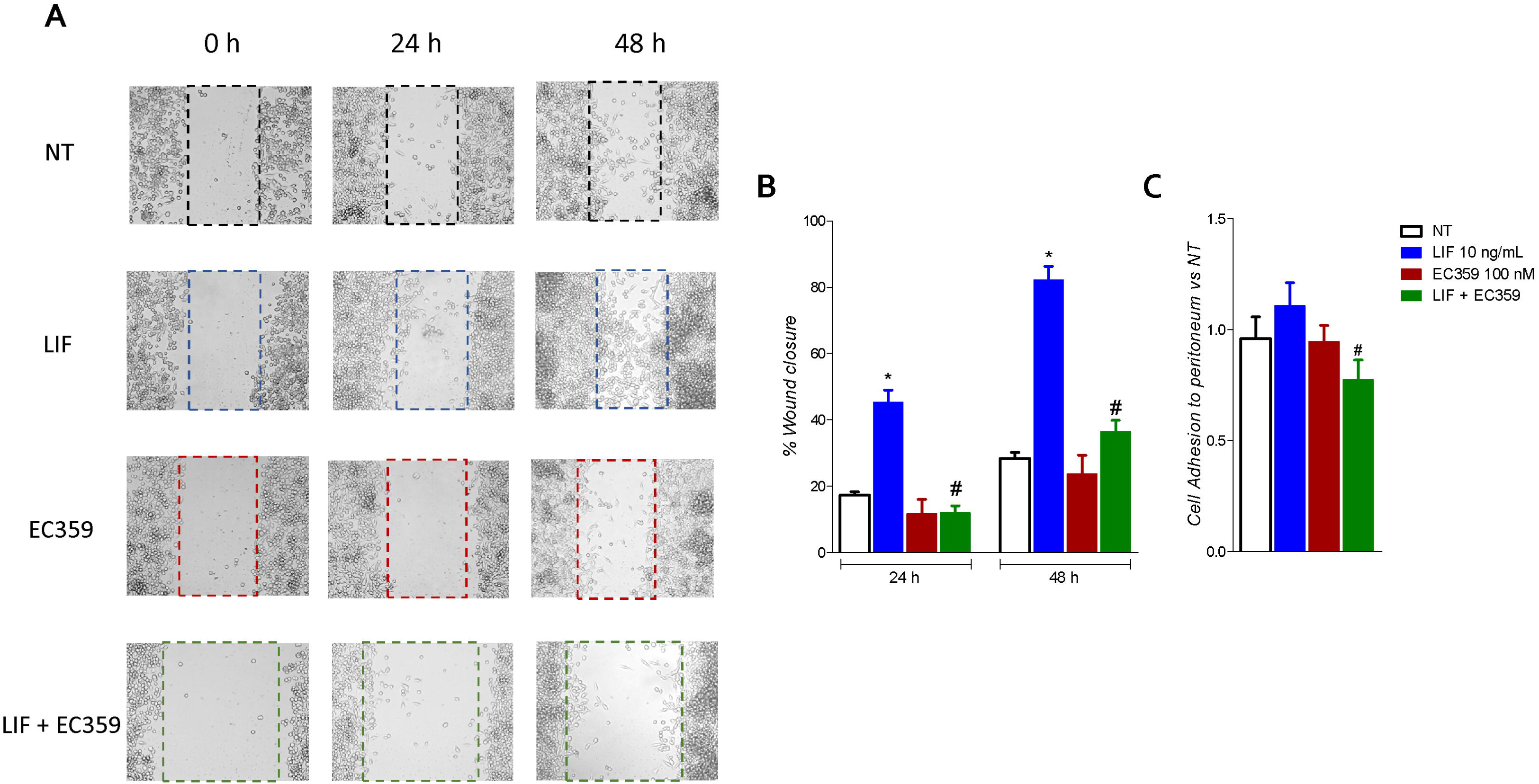
LIFR antagonism inhibits MKN45. **A)** A scratch wound healing assay is shown: MKN45 cell monolayers were scraped in a straight line using a p200 pipette tip in order to create a “scratch”, then they are left untreated or primed with LIF 10 ng/ml, EC359 100 nM and LIF + EC359. The wound generated was imaged at 0, 24 and 48 hours of incubation with the indicated compounds. The images show cell migration at the three times point indicated. Images obtained points were analyzed measuring scraped area and its closure versus the first time point, 0 hours. Results are the mean ± SEM of 3 samples per group. **B)** Cell adhesion to peritoneum. Experiment was conducted in quintuplicate. (* represents statistical significance versus NT, and # versus LIF, p < 0.05).

In summary, results presented manifest that LIFR inhibition decrease the LIF-induced ability to gain the migratory phenotype of MKN45 cells.

### Discussion and conclusion

The LIF/LIFR signaling has been identified as a potential therapeutic target in the treatment of several cancers. In the present study, we report the transcriptome profiling (RNAseq) of a group of patients with GC with or without peritoneal carcinomatosis and the identification of LIFR as one of the highest expressed genes in patients with peritoneal involvement and as a strong predictor of a poor prognosis in these patients. In addition, we have shown that activation of LIF/LIFR signalling in GC cells promotes the acquisition of a mesenchymal phenotype suggesting a potential mechanistic role of LIF/LIFR signalling in development of peritoneal metastasis.

The peritoneal carcinomatosis is a relatively common localization of metastasis in gastric cancer, occurring in up to 14% of newly diagnosed gastric cancer patients, and is the most common site (∼50%) of recurrence in gastric cancer patients after radical surgery (41–43). In our series, mean survival time, of patients with positive peritoneal cytology or macroscopic evidence of peritoneal metastasis at surgery (Cy+/P+) was approximately 12 months and was significantly lower in comparison to the 60 months median survival observed in patients that were Cy0/T0. These data are in agreement with previous findings, confirming the fact that the development of a peritoneal disease is a strong predictor of poor prognosis in GC patients.

In addition to the presence of peritoneal disease, by the transcriptome in paired samples of neoplastic tissue and normal mucosa, we have identified a group of 6 differentially expressed gene, that predict poor prognosis: OGN, FABP1, LIF, ONECUT2, SFRP2 and GPA33. More specifically we have shown that the top three upregulated genes ONG, LIFR and SFRP2 and the top three downregulated genes, FABP1, ONECUT2 and GPA33 associates with a poor prognosis, but statistically significantly difference were detected only for the expression of ONECUT2 and LIFR (P< 0.05).

ONECUT2 belong to the family of the ONECUT transcription factors, a small group are evolutionarily conserved proteins that play a role in the development of the liver, pancreas and neuronal system (44). Although a role for ONECUT2 in cancer is not well defined, it has been demonstrated that the expression of this gene is aberrantly upregulated in a variety of cancers including hepatocellular carcinoma, prostate cancer, colorectal cancer and ovarian cancer, suggesting a role for this transcription factor in the modulation of cancer progression (45). Despite the fact that, similar to a previous study, we have found that ONECUT2 expression was increased in the neoplastic tissues in comparison to paired samples obtained from non-neoplastic mucosa, but we have found that reduced levels of ONECUT2, rather than its induction, is a poor prognosis predictor in patients with GC and PC (39). The reason for this discrepancy is unclear, because overexpression of ONECUT2 in MKN54 and AGS, two gastric cancer cell lines, promotes cell proliferation and migration. However, others have reported that ONECUT2 regulation occurs through epigenetic regulation and hypomethylation of that CpGs in the promoter of ONECUT2 occurs primarily in intestinal metaplasia and gastric cancers, and that hypomethylation was associated with ONECUT2 upregulation. It is therefore suggested that that ONECUT2 might promote intestinal differentiation or development of gastric cancer and could be used as a early detection biomarkers of gastric carcinogenesis, while its role in advanced disease is less defined (46).

The metastatic cascade is a multistep process whereby cancer cells detach from primary tumor, migrate and attach to distant peritoneum, followed by invasion into sub- peritoneal tissues and cell proliferation to form detectable metastasis (47). Despite the clinical relevance, however, the specific molecular mechanisms that drive the formation of peritoneal metastasis in GC remain poorly understood, although previous studies using paired samples of primary and metastatic tumors have identified several putative mediators, mostly involved in EMT remodeling, cell motility and cytoskeleton rearrangement (48). Here we report, that development of peritoneal disease in our series of patients with GC, associates with a robust upregulation of the LIFR in the primary tumors. This finding prompted us to further investigate whether the LIF/LIFR system was involved in promoting the EMT phenotype, a process that involves a deep reprogramming of the cancer cells genes. LIFR is an heterodimeric membrane receptor complex composed by LIFRβ and GP130 (49) (50) and while the receptor lacks an intrinsic tyrosine kinase activity, the LIFR/GP130 complex constitutively associates with JAK-Tyk family of cytoplasmic tyrosine kinases, which facilitates downstream signaling and STAT3 phosphorylation activation. Several tumors exhibit upregulated JAK/STAT, ERK/MAPK and PI3K/AKT signaling via autocrine or paracrine activation of LIF/LIFR GP130 and this pathway significantly contribute to EMT in several cancers, disease progression and a poorer relapse-free survival in several cancers (51). LIF also participate to cross-talk between tumor cells and matrix fibroblasts to mediate the pro-invasive activation of stromal fibroblasts (52) and promotes drug resistance to HDAC inhibitors (53). Here, by using GC cell lines we have shown that LIFR expression varies from one line to another, and that MKN45 were the cells with the highest expression. Challenging these cells with LIF promotes the acquisition of migratory phenotype, and this associates with acquisition of molecular signature of EMT and these changes associate with LIF/LIFR signaling as assessed by measuring JAK and STAT3 phosphorylation. Of relevance, the LIFR inhibitor, EC359 (IC_50_ values of 10.2 nM) reversed these changed. Cotreating MKN45 cells with EC359, reversed changes promoted by LIF on cell and JAK and STAT3 phosphorylation in vitro as well as the regulation of E-Cadherin and vimentin in a dose-dependent manner, further confirming the role of LIF/LIFR in supporting migration and invasion of GC cells.

Cytoskeletal remodeling is closely correlated with tumor migration, invasion and metastasis (54). LIFR plays an essential role in regulation of actin filament dynamics by modulating the expression of Vimentin. Consistent with this background we demonstrated that pharmacological inhibition of LIFR negatively regulated the expression of vimentin and that this effects associated with a reduce cell motility and impaired migration (55) (56). Vimentin plays an important role in tumor invasion and metastasis (57), and its counter-regulation is a further evidence of the role the LIF/LIFR signaling plays in the modulation of EMT process. Of relevance, LIFR activation positively regulates vimentin expression and downregulates E-cadherin vi JAK and STAT3 phosphorylation, LIFR antagonism reversed this pathway (55) (56) (57).

In conclusion, by NGS RNAseq analysis we have identified a the LIF/LIFR pathway as an important mechanism in disease progression in GC. High levels of expression of LIFR mRNA predict poor prognosis a reduced response to therapy. Additionally, by using GC cell lines we have shown that LIFR activation results in JAK STAT3 phosphorylation, and EMT as demonstrated by vimentin induction and blunted expression of E-Cadherin. These molecular changes associate with a migratory phenotype of GC cell lines and reversed by LIF/LIFR antagonism. Together we suggest that targeting LIF/LIFR signaling might have utility in management of GC.

## Authors contributions

SF, LG, and AD contributed to conception and design of the study. AZ and SF provided research funding. LG, EM and AD provided human samples. CD, SM, EM, and MB performed the data analysis. CD, SM and EM performed the statistical analysis. SF, CD, SM and EM wrote the manuscript. CD, SM, RR, MB, RB and UG contributed experimental settings. All authors contributed to manuscript revision, read, and approved the submitted version.

---

Supplementary Figure 1. MKN45 cells were serum starved and left untreated or primed with LIF (10,50,100 ng/ml). Data shown are: **A)** MTS assay. Each value is expressed relative to those of non-treated (NT), which are arbitrarily settled to 1. Results are the mean ± SEM of 10 samples per group. Relative mRNA expression of **B)** the proliferation marker C-MYC; and EMT markers: E-CADHERIN; SNAL-1; and VIMENTIN. Each value is normalized to GAPDH and is expressed relative to those of NT, which are arbitrarily settled to 1. Results are the mean ± SEM of 5 samples per group. (* represents statistical significance versus NT, and # versus LIF, p < 0.05). Cell cycle phases analysis were performed by Ki-67/DAPI staining through IC-FACS analysis. Data shown are: percentage of **D)** Representative IC- FACS showed cell cycle fraction and apoptosis rate in NT, LIF 10 ng/ml, EC359 25 nM and LIF plus EC359. **E)** cell in G0-G1 cell cycle phases, S-G2-M cell cycle phases and ratio between % G0-G1 and % S-G2-M; **F)** Percentage of Apoptotic cells. Results are the mean ± SEM of 3 samples per group. (* represents statistical significance versus NT, and # versus LIF, p < 0.05).

## References

1. Torre LA, Bray F, Siegel RL, Ferlay J, Lortet-Tieulent J, Jemal A. Global cancer statistics, 2012. CA Cancer J Clin [Internet]. 2015/02/04. 2015;65(2):87–108. Available from: https://www.ncbi.nlm.nih.gov/pubmed/25651787

2. Bray F, Ferlay J, Soerjomataram I, Siegel RL, Torre LA, Jemal A. Global cancer statistics 2018: GLOBOCAN estimates of incidence and mortality worldwide for 36 cancers in 185 countries. CA Cancer J Clin [Internet]. 2018/09/12. 2018;68(6):394–424. Available from: https://www.ncbi.nlm.nih.gov/pubmed/30207593

3. Ahmad SA, Xia BT, Bailey CE, Abbott DE, Helmink BA, Daly MC, et al. An update on gastric cancer. Curr Probl Surg. 2016 Oct;53(10):449–90.

4. González CA, Sala N, Rokkas T. Gastric cancer: epidemiologic aspects. Helicobacter. 2013/09/18. 2013 Sep;18 Suppl 1:34–8.

5. Di Giorgio C, Roselli R, Biagioli M, Marchianò S, Distrutti E, Bordoni M, et al. Organoids as ex vivo culture system to investigate infection-host interaction in gastric pre-carcinogenesis. Recent Adv Inflamm allergy drug Discov. 2022 Jan;

6. Lauren P. THE TWO HISTOLOGICAL MAIN TYPES OF GASTRIC CARCINOMA: DIFFUSE AND SO-CALLED INTESTINAL-TYPE CARCINOMA. AN ATTEMPT AT A HISTO-CLINICAL CLASSIFICATION. Acta Pathol Microbiol Scand [Internet]. 1965/01/01. 1965;64:31–49. Available from: https://www.ncbi.nlm.nih.gov/pubmed/14320675

7. Carino A, Graziosi L, Marchianò S, Biagioli M, Marino E, Sepe V, et al. Analysis of Gastric Cancer Transcriptome Allows the Identification of Histotype Specific Molecular Signatures With Prognostic Potential. Front Oncol. 2021;11:663771.

8. Chia N-Y, Tan P. Molecular classification of gastric cancer. Ann Oncol Off J Eur Soc Med Oncol. 2016 May;27(5):763–9.

9. Tan P, Yeoh K-GG. Genetics and Molecular Pathogenesis of Gastric Adenocarcinoma. Gastroenterology. 2015/06/16. 2015 Oct;149(5):1153–1162.e3.

10. Lutz MP, Zalcberg JR, Ducreux M, Adenis A, Allum W, Aust D, et al. The 4th St. Gallen EORTC Gastrointestinal Cancer Conference: Controversial issues in the multimodal primary treatment of gastric, junctional and oesophageal adenocarcinoma. Eur J Cancer. 2019 May;112:1–8.

11. Lei Z, Tan IB, Das K, Deng N, Zouridis H, Pattison S, et al. Identification of molecular subtypes of gastric cancer with different responses to PI3-kinase inhibitors and 5- fluorouracil. Gastroenterology [Internet]. 2013/05/14. 2013;145(3):554–65. Available from: https://www.ncbi.nlm.nih.gov/pubmed/23684942

12. Network CGAR. Comprehensive molecular characterization of gastric adenocarcinoma. Nature [Internet]. 2014/08/01. 2014 Sep;513(7517):202–9. Available from: https://www.ncbi.nlm.nih.gov/pubmed/25079317

13. Cristescu R, Lee JH, Nebozhyn M, Kim KM, Ting JC, Wong SS, et al. Molecular analysis of gastric cancer identifies subtypes associated with distinct clinical outcomes. Nat Med [Internet]. 2015/04/20. 2015;21(5):449–56. Available from: https://www.ncbi.nlm.nih.gov/pubmed/25894828

14. Ajani JA, Lee J, Sano T, Janjigian YY, Fan D, Song S. Gastric adenocarcinoma. Nat Rev Dis Prim. 2017 Jun;3:17036.

15. Schmidt B, Yoon SS. D1 versus D2 lymphadenectomy for gastric cancer. J Surg Oncol. 2013 Mar;107(3):259–64.

16. Peinado H, Zhang H, Matei IR, Costa-Silva B, Hoshino A, Rodrigues G, et al. Pre-metastatic niches: organ-specific homes for metastases. Nat Rev Cancer. 2017 May;17(5):302–17.

17. Liu D, Li C, Trojanowicz B, Li X, Shi D, Zhan C, et al. CD97 promotion of gastric carcinoma lymphatic metastasis is exosome dependent. Gastric cancer Off J Int Gastric Cancer Assoc Japanese Gastric Cancer Assoc. 2016 Jul;19(3):754–66.

18. Kim MA, Lee HS, Lee HE, Kim JH, Yang H-K, Kim WH. Prognostic importance of epithelial-mesenchymal transition-related protein expression in gastric carcinoma. Histopathology. 2009 Mar;54(4):442–51.

19. Ye X, Weinberg RA. Epithelial-Mesenchymal Plasticity: A Central Regulator of Cancer Progression. Trends Cell Biol. 2015 Nov;25(11):675–86.

20. Marano L, Marrelli D, Sammartino P, Biacchi D, Graziosi L, Marino E, et al. Cytoreductive Surgery and Hyperthermic Intraperitoneal Chemotherapy for Gastric Cancer with Synchronous Peritoneal Metastases: Multicenter Study of “Italian Peritoneal Surface Malignancies Oncoteam-S.I.C.O.”. Ann Surg Oncol. 2021 Dec;28(13):9060–70.

21. Thomassen I, Bernards N, van Gestel YR, Creemers G-J, Jacobs EM, Lemmens VE, et al. Chemotherapy as palliative treatment for peritoneal carcinomatosis of gastric origin. Vol. 53, Acta oncologica (Stockholm, Sweden). England; 2014. p. 429–32.

22. Marano L, Polom K, Patriti A, Roviello G, Falco G, Stracqualursi A, et al. Surgical management of advanced gastric cancer: An evolving issue. Eur J Surg Oncol [Internet]. 2015/11/14. 2016;42(1):18–27. Available from: https://www.ncbi.nlm.nih.gov/pubmed/26632080

23. Shiozaki H, Elimova E, Slack RS, Chen HC, Staerkel GA, Sneige N, et al. Prognosis of gastric adenocarcinoma patients with various burdens of peritoneal metastases. J Surg Oncol [Internet]. 2015/11/25. 2016;113(1):29–35. Available from: https://www.ncbi.nlm.nih.gov/pubmed/26603684

24. Seeneevassen L, Martin OCB, Lehours P, Dubus P, Varon C. Leukaemia inhibitory factor in gastric cancer: friend or foe? Gastric cancer Off J Int Gastric Cancer Assoc Japanese Gastric Cancer Assoc. 2022 Mar;25(2):299–305.

25. Pinho V, Fernandes M, da Costa A, Machado R, Gomes AC. Leukemia inhibitory factor: Recent advances and implications in biotechnology. Cytokine Growth Factor Rev. 2020 Apr;52:25–33.

26. Wang M-T, Fer N, Galeas J, Collisson EA, Kim SE, Sharib J, et al. Blockade of leukemia inhibitory factor as a therapeutic approach to KRAS driven pancreatic cancer. Nat Commun. 2019 Jul;10(1):3055.

27. Yu H, Yue X, Zhao Y, Li X, Wu L, Zhang C, et al. LIF negatively regulates tumour-suppressor p53 through Stat3/ID1/MDM2 in colorectal cancers. Nat Commun. 2014 Oct;5:5218.

28. Wang J, Xie C, Pan S, Liang Y, Han J, Lan Y, et al. N-myc downstream-regulated gene 2 inhibits human cholangiocarcinoma progression and is regulated by leukemia inhibitory factor/MicroRNA-181c negative feedback pathway. Hepatology. 2016 Nov;64(5):1606–22.

29. Li X, Yang Q, Yu H, Wu L, Zhao Y, Zhang C, et al. LIF promotes tumorigenesis and metastasis of breast cancer through the AKT-mTOR pathway. Oncotarget. 2014 Feb;5(3):788–801.

30. Nicola NA, Babon JJ. Leukemia inhibitory factor (LIF). Cytokine Growth Factor Rev. 2015 Oct;26(5):533–44.

31. Viswanadhapalli S, Luo Y, Sareddy GR, Santhamma B, Zhou M, Li M, et al. EC359: A First-in-Class Small-Molecule Inhibitor for Targeting Oncogenic LIFR Signaling in Triple-Negative Breast Cancer. Mol Cancer Ther. 2019 Aug;18(8):1341–54.

32. Zhao X, Ye F, Chen L, Lu W, Xie X. Human epithelial ovarian carcinoma cell-derived cytokines cooperatively induce activated CD4+CD25-CD45RA+ naïve T cells to express forkhead box protein 3 and exhibit suppressive ability in vitro. Cancer Sci. 2009 Nov;100(11):2143–51.

33. Pascual-García M, Bonfill-Teixidor E, Planas-Rigol E, Rubio-Perez C, Iurlaro R, Arias A, et al. LIF regulates CXCL9 in tumor-associated macrophages and prevents CD8(+) T cell tumor-infiltration impairing anti-PD1 therapy. Nat Commun. 2019 Jun;10(1):2416.

34. Lin S-R, Wen Y-C, Yeh H-L, Jiang K-C, Chen W-H, Mokgautsi N, et al. EGFR- upregulated LIFR promotes SUCLG2-dependent castration resistance and neuroendocrine differentiation of prostate cancer. Oncogene. 2020 Oct;39(44):6757–75.

35. Bian S-B, Yang Y, Liang W-Q, Zhang K-C, Chen L, Zhang Z-T. Leukemia inhibitory factor promotes gastric cancer cell proliferation, migration, and invasion via the LIFR- Hippo-YAP pathway. Ann N Y Acad Sci. 2021 Jan;1484(1):74–89.

36. Seeneevassen L, Giraud J, Molina-Castro S, Sifré E, Tiffon C, Beauvoit C, et al. Leukaemia Inhibitory Factor (LIF) Inhibits Cancer Stem Cells Tumorigenic Properties through Hippo Kinases Activation in Gastric Cancer. Cancers (Basel). 2020 Jul;12(8).

37. Chen Y. Scratch Wound Healing Assay. Bio-protocol [Internet]. 2012;2(5):e100. Available from: https://doi.org/10.21769/BioProtoc.100

38. Asao T, Yazawa S, Kudo S, Takenoshita S, Nagamachi Y. A novel ex vivo method for assaying adhesion of cancer cells to the peritoneum. Cancer Lett. 1994 Apr;78(1– 3):57–62.

39. Chen J, Chen J, Sun B, Wu J, Du C. ONECUT2 Accelerates Tumor Proliferation Through Activating ROCK1 Expression in Gastric Cancer. Cancer Manag Res. 2020;12:6113–21.

40. Huang L, Wu R-L, Xu A-M. Epithelial-mesenchymal transition in gastric cancer. Am J Transl Res. 2015;7(11):2141–58.

41. Thomassen I, van Gestel YR, van Ramshorst B, Luyer MD, Bosscha K, Nienhuijs SW, et al. Peritoneal carcinomatosis of gastric origin: a population-based study on incidence, survival and risk factors. Int J cancer. 2014 Feb;134(3):622–8.

42. Sugarbaker PH, Yu W, Yonemura Y. Gastrectomy, peritonectomy, and perioperative intraperitoneal chemotherapy: the evolution of treatment strategies for advanced gastric cancer. Semin Surg Oncol. 2003;21(4):233–48.

43. Wadhwa R, Song S, Lee J-S, Yao Y, Wei Q, Ajani JA. Gastric cancer-molecular and clinical dimensions. Nat Rev Clin Oncol. 2013 Nov;10(11):643–55.

44. Vanhorenbeeck V, Jenny M, Cornut J-F, Gradwohl G, Lemaigre FP, Rousseau GG, et al. Role of the Onecut transcription factors in pancreas morphogenesis and in pancreatic and enteric endocrine differentiation. Dev Biol. 2007 May;305(2):685–94.

45. Yu J, Li D, Jiang H. Emerging role of ONECUT2 in tumors. Oncol Lett. 2020 Dec;20(6):328.

46. Seo E-H, Kim H-J, Kim J-H, Lim B, Park J-L, Kim S-Y, et al. ONECUT2 upregulation is associated with CpG hypomethylation at promoter-proximal DNA in gastric cancer and triggers ACSL5. Int J cancer. 2020 Jun;146(12):3354–68.

47. Sun F, Feng M, Guan W. Mechanisms of peritoneal dissemination in gastric cancer. Oncol Lett. 2017 Dec;14(6):6991–8.

48. Kang X, Li W, Liu W, Liang H, Deng J, Wong CC, et al. LIMK1 promotes peritoneal metastasis of gastric cancer and is a therapeutic target. Oncogene. 2021 May;40(19):3422–33.

49. Metcalf D. The unsolved enigmas of leukemia inhibitory factor. Stem Cells. 2003;21(1):5–14.

50. Stahl N, Boulton TG, Farruggella T, Ip NY, Davis S, Witthuhn BA, et al. Association and activation of Jak-Tyk kinases by CNTF-LIF-OSM-IL-6 beta receptor components. Science. 1994 Jan;263(5143):92–5.

51. Yue X, Zhao Y, Zhang C, Li J, Liu Z, Liu J, et al. Leukemia inhibitory factor promotes EMT through STAT3-dependent miR-21 induction. Oncotarget. 2016 Jan;7(4):3777–90.

52. Albrengues J, Bourget I, Pons C, Butet V, Hofman P, Tartare-Deckert S, et al. LIF mediates proinvasive activation of stromal fibroblasts in cancer. Cell Rep. 2014 Jun;7(5):1664–78.

53. Zeng H, Qu J, Jin N, Xu J, Lin C, Chen Y, et al. Feedback Activation of Leukemia Inhibitory Factor Receptor Limits Response to Histone Deacetylase Inhibitors in Breast Cancer. Cancer Cell. 2016 Sep;30(3):459–73.

54. Fife CM, McCarroll JA, Kavallaris M. Movers and shakers: cell cytoskeleton in cancer metastasis. Br J Pharmacol. 2014 Dec;171(24):5507–23.

55. Mei J-W, Yang Z-Y, Xiang H-G, Bao R, Ye Y-Y, Ren T, et al. MicroRNA-1275 inhibits cell migration and invasion in gastric cancer by regulating vimentin and E-cadherin via JAZF1. BMC Cancer. 2019 Jul;19(1):740.

56. Ueda T, Volinia S, Okumura H, Shimizu M, Taccioli C, Rossi S, et al. Relation between microRNA expression and progression and prognosis of gastric cancer: a microRNA expression analysis. Lancet Oncol. 2010 Feb;11(2):136–46.

57. Du L, Li J, Lei L, He H, Chen E, Dong J, et al. High Vimentin Expression Predicts a Poor Prognosis and Progression in Colorectal Cancer: A Study with Meta-Analysis and TCGA Database. Biomed Res Int. 2018;2018:6387810.

