## Supplementary figures and images for "Next generation sequencing analysis of gastric cancer identifies the leukemia inhibitory factor receptor (LIFR) as a driving factor in gastric cancer progression and as a predictor of poor prognosis"

### Figure S1.tif

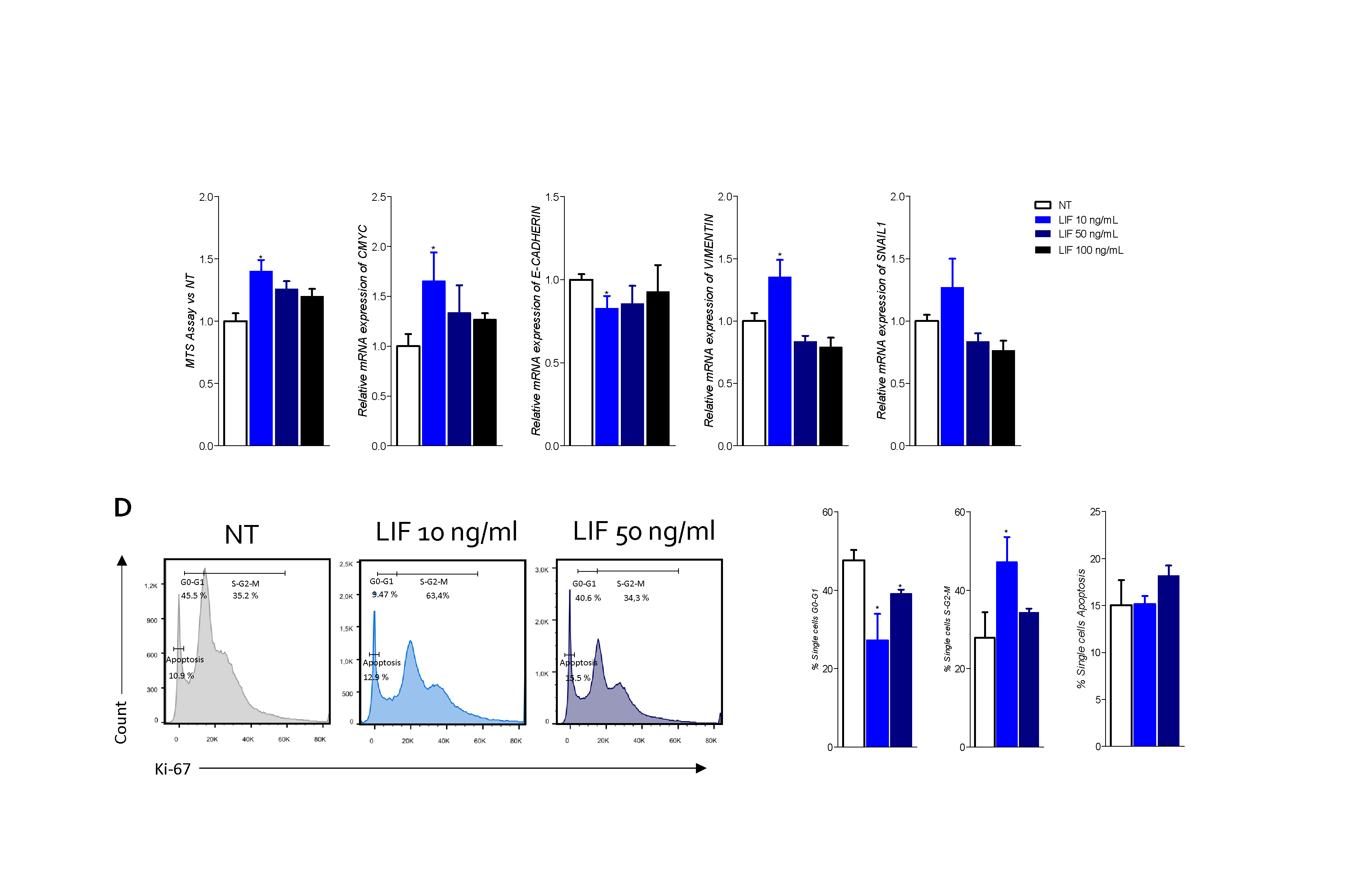
